# A genome-wide CRISPR/Cas phenotypic screen for modulators of DUX4 cytotoxicity reveals screen complications

**DOI:** 10.1101/2020.07.27.223420

**Authors:** Ator Ashoti, Francesco Limone, Melissa van Kranenburg, Anna Alemany, Mirna Baak, Judith Vivié, Federica Piccioni, Menno Creyghton, Kevin Eggan, Niels Geijsen

**Affiliations:** Hubrecht Institute, Developmental Biology and Stem Cell Research, Utrecht, Netherlands; Department of Stem Cell and Regenerative Biology, Harvard University Cambridge, MA, USA; Stanley Center for Psychiatric Research, Broad Institute of MIT and Harvard, Cambridge, MA, USA; Single Cell Discoveries, Utrecht, Netherlands; Leiden University Medical Center, Leiden, The Netherlands; Erasmus University Medical Centre, Rotterdam, The Netherlands; Broad institute, Cambridge, MA, USA

## Abstract

Facioscapulohumeral muscular dystrophy (FHSD), a fundamentally complex muscle disorder that thus far remains untreatable. As the name implies, FSHD starts in the muscles of the face and shoulder gridle. The main perturbator of the disease is the pioneer transcription factor DUX4, which is misexpressed in affected tissues due to a failure in epigenetic repressive mechanisms. In pursuit of unraveling the underlying mechanism of FSHD and finding potential therapeutic targets or treatment options, we performed an exhaustive genome-wide CRISPR/Cas9 phenotypic rescue screen to identify modulators of DUX4 cytotoxicity. We found no key effectors other than DUX4 itself, suggesting treatment efforts in FSHD should be directed towards its direct modulation.

The screen did however reveal some rare and unexpected Cas9-induced genomic events, that may provide important considerations for planning future CRISPR/Cas9 knock-out screens.

## Introduction

Facioscapulohumeral muscular dystrophy (FSHD) is an autosomal dominant degenerative muscle disease. It’s one of the most prevalent neuromuscular disorder^1^, characterized by progressive and asymmetric muscle weakness which generally starts in facial muscles, and then slowly progresses to muscles of the shoulders, upper limbs and eventually the lower extremities^2^. Age of onset is highly variable, but calculations based on a 122 case study demonstrates that the mean age of onset is in the early twenties (21-23)^3^. The primary cause of the disease is the misexpression of the double homeobox 4 (DUX4) transcription factor, due to failure in epigenetic silencing^3–6^. DUX4 is normally expressed early in development in the cleavage stage embryo^7,8^, in the adult testis^6^ and in the thymus^9^. De-repression of DUX4 in muscle activates a large cascade of events, triggering the activation of many pathways^8,10–19^, with target genes being involved in biological processes such as RNA splicing and processing (DBR1^10,20–22^, CWC15^10,20,22^, PNN^10,21^, CLP1^10,21,22^, TFIP11^10,20–22^), spermatogenesis (CCNA1^10,20–22^, ZNF296^10,20–22^, TESK2^10,20,21^), early embryonic development (ZSCAN4^10,20–22^, LEUTX^20–22^, STIL^10,20,21^), protein processing and degradation (SIAH1^10,20–22^, RHOBTB1^10,20,21^, TRIM36^10,20,21^), and cell motility and migration (CXCR4^10,20,21^, ROCK1^10,21^, SNAI1^10,20–22^).

We hypothesized that of one or more factors downstream of DUX4 expression are responsible for the rapid apoptotic response that follows DUX4 induction. Knowing if there are key downstream targets of DUX4 can have important clinical applications as they could direct intelligent therapeutic design. We tested this hypothesis by performing a genome wide CRISPR/Cas9 knockout screen.

CRISPR/Cas9 which is now a highly popular and widely used genome editing technique, was initially discovered as the adaptive immune system of bacteria, to protect against viral infection^23,24^. Although not the first genome editing method, CRISPR/Cas9 has proven to be much more user friendly due to its easy manipulability, and being more cost-, labor- and time-efficient compared to its predecessors: transcription activator-like effector nucleases (TALENs)^25–28^ and ZINC-fingers nucleases (ZFNs)^29–33^. Its ability to knock-out any gene by creating a double stranded break^34–36^ in such an easy manner, makes this technique very suitable for genome-wide loss off function studies. The advantages and ease-of-use of the CRISPR/Cas9 technology inspired us to perform a genome wide screen on a FSHD *in-vitro* model, to find potential modulators that contribute or aggravate the FSHD pathophysiology. Successful performance of a FSHD genome-wide screen will critically depend on the cell system being used. The cells should be highly proliferative, easily transfected and display a robust DUX4-induced phenotype. Fortunately, DUX4 is a so-called pioneer factor^49,50^, capable of regulating its target genes independent of their chromatin-state. The network of genes activated by pioneer factors is therefore less affected by cellular identity. Indeed, Jones and colleagues have demonstrated that DUX4 activates the same downstream target genes in B-lymphocytes as previously identified in skeletal muscle myoblasts^51,52^. Using an adherent leukemic cell line that is frequently used for genome-wide screening purposes (HAP1^53,54^), we performed an exhaustive CRISPR knockout screen to identify factors that could mitigate DUX4-induced cytotoxicity. We inserted a doxycycline-inducible DUX4 transgene into the HAP1 cells ^53,54^ to generate DUX4 inducible expression (DIE) cells. Using the Brunello CRISPR/Cas9 library^55^, we screened for modulators of DUX4 cytotoxicity. Our results suggest that no single gene knockout is capable of rescuing DUX4-triggered apoptosis in our transgene model system.

This study does however, provide some interesting insight into critical parameters that need to be considered when executing a genome-wide CRISPR screen.

## Results

### Generation and validation of a DUX4 inducible cell line

In order to perform a successful genome-wide screen, a cellular model is required that is highly proliferative, easily transfected and which displays a robust phenotype. HAP1 cells display all these characteristics and have been used extensively in a wide variety of functional screens^54,56–64^. The cells were initially reported to be near-haploid^53,54^, but subsequently rediploidized^65,66^. These rediploidized cells were used to generate the inducible DUX4 cell line. Since even low levels of DUX4 expression can efficiently induce apoptosis ^6,20,67^, we needed to circumvent premature DUX4 toxicity caused by leaky expression of the Tet-On system^68,69^ during the generation of the cell line. To avoid this, we inserted a LoxP-DsRed-LoxP stop-cassette in between the Tet-operator and a DUX4 transgene. The DUX4 transgene element itself consisted of the three exons and the two introns of *DUX4* including the polyA terminal sequence (haplotype 4A161^70^). After stable integration of the construct, the stop-cassette was removed using CRE recombinase, thereby placing *DUX4* under the control of the Tet operator (Fig. 1A). 80 monoclonal lines were derived by single cell flow-cytometry sorting, of which 1 demonstrated DUX4 induction and robust cell death upon doxycycline addition (Fig. 1B), which we named ‘DUX4-inducible expression’ (DIE) cells.

**Figure 1.**
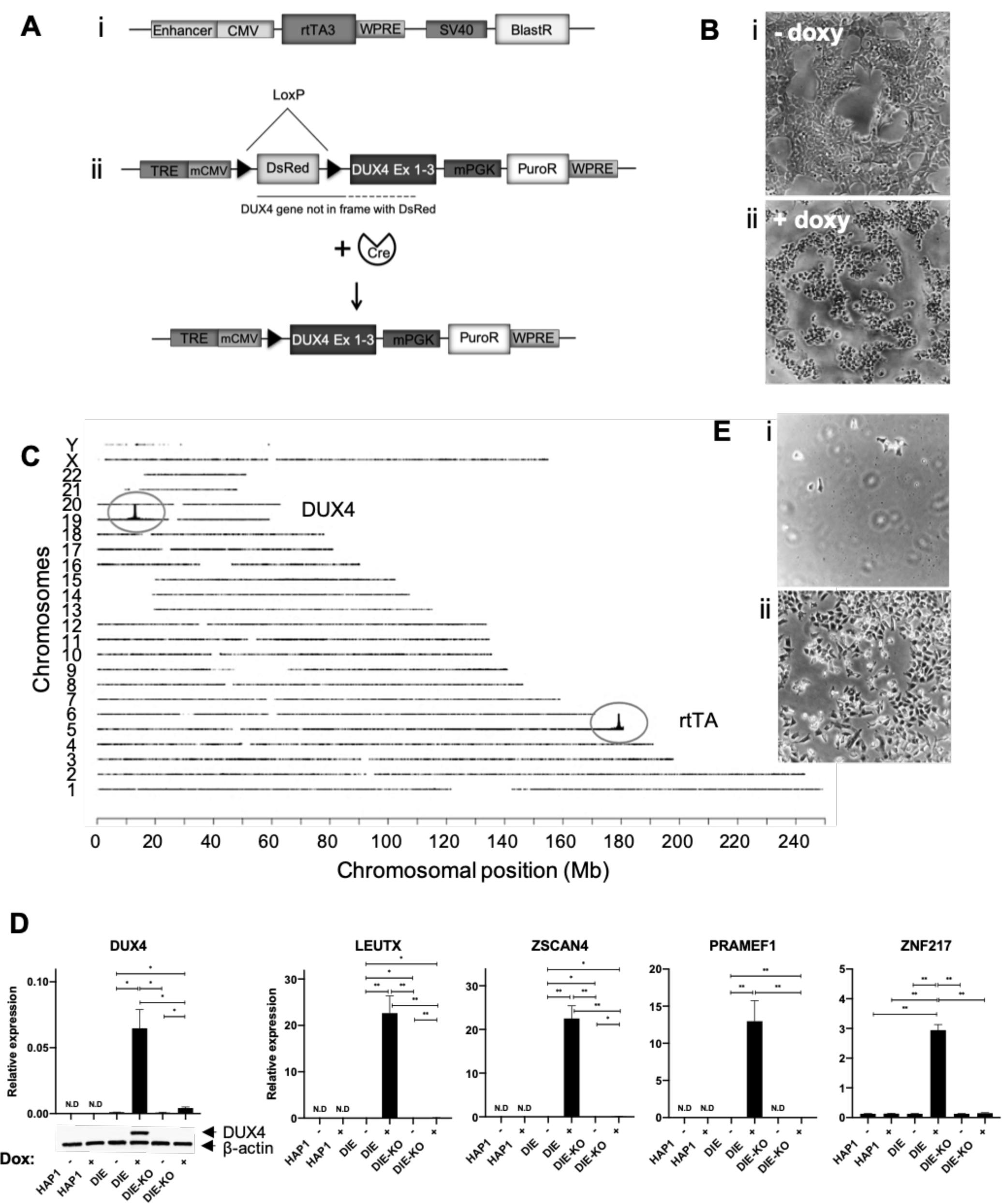
Creation and validation of the DIE cell line. **(A)** Schematic representation of the i) rtTA construct, ii) the inducible LSL-DUX4 cassettes, and the inducible DUX4 cassette upon removal of the stop cassette (LSL) through CRE recombinase. **(B)** Phase contrast images of DIE cells i) without doxycycline exposure and ii) with 24h of doxycycline exposure. **(C)** Schematic representation of transgene integration sites within human genome, by TLA analysis. The inducible DUX4 cassette maps back to the p-arm of chromosome 19, and the rtTA transgene maps back to the end of the q-arm of chromosome 5. **(D)** Expression of DUX4 mRNA and protein in HAP1, DIE and DIE-KO cells with or without doxycycline admission, as detected by qRT-PCR and western blot analysis. β -actin served as a loading control in the western blot. **(E)** Phase contrast images of doxycycline exposed DIE cells which were transduced with either i) only Cas9 protein, or ii) Cas9 protein and DUX4 sgRNA. Dead cells were removed by a DPBS wash to expose the surviving population. **(F)** mRNA expression of known downstream targets of DUX4 in HAP1, DIE and DIE-KO cells with or without doxycycline admission, as measured by qRT-PCR.

To identify the sites of integration of the rtTA/BlastR and DUX4/PuroR transgenes in the DIE cells, we performed targeted locus amplification (TLA)^71^. TLA analysis showed a single integration site for both the rtTA and DUX4 transgenes (Fig. 1C). The DUX4 cassette integrated in the p-arm of chromosome 19, within the MAST1 gene, and the rtTA cassette integrated in the MGAT4B gene which is located at the end of chromosome 5q.

To further analyze the functional effect of DUX4 induction, DIE cells were stained with AnnexinV-Alexa Fluor 488 and propidium iodine (PI) (Fig. S2). As shown in the supplementary video’s, DIE cells stained positive for the apoptotic marker AnnexinV, during 12 hours of doxycycline exposure. To show that the apoptotic phenotype was dependent on the induction of the DUX4 transgene, we targeted the DUX4 transgene using CRISPR/Cas9 and 2 independent guide RNAs (gRNAs), one targeting the C-terminal domain of DUX4 and the other close to the polyA tail of DUX4. RT-qPCR and Western blot (WB) analysis of the DIE and the DIE-DUX4 knockout (DIE-KO) cells demonstrated successful knockout of the DUX4 transgene at RNA and protein level (Fig. 1D). In addition, CRISPR/Cas9 targeting of the DUX4 transgene successfully rescued the DIE cells from apoptosis upon doxycycline administration, demonstrating that apoptosis upon doxycycline induction in the DIE line is mediated by DUX4 (Fig. 1E). DUX4 induction in the DIE cells also resulted in induction of its known downstream target genes (LEUTX, ZSCAN4, PRAMFE1 and ZNF217) (Fig. 1F), demonstrating that inducing DUX4 expression induces downstream target genes that are also induced by endogenous DUX4.

### DUX4 gene expression signature in DIE cells

Next we analyzed the downstream transcriptional changes that were induced by DUX4 in the DIE cells with RNA sequencing. We compared 4 induced and uninduced technical replicates of two lines; the DIE line, and the DIE-KO line. As shown in Fig. 2A, DUX4 induction resulted in progressive temporal changes in gene expression. To visualize the magnitude and speed of the DUX4-induced transcriptional changes, we plotted the log2fold change of all detected transcripts against the density of differentially expressed genes at any given log2fold change (Fig. 2B). As shown, DUX4 induction results in a profound and progressive increase in the number of differentially expressed genes; with 64 differentially expressed transcripts at 4.5 hours post DUX4 induction and 467 differentially expressed transcripts at 8.5 hours (Fig. 2B and 2C). The number of genes of which the expression is induced is greater than those with reduced expression levels, consistent with DUX4’s role as an activating pioneering factor^49^(Fig. 2B). This is further reflected by the differential expression analysis, demonstrating an increase in upregulated genes in both induced DIE samples compared to uninduced DIE sample [Padj value ≤ 0.01, Log2foldchange ≥ 1] (Fig. 2C and D, Supplementary Table S1). Most differentially expressed genes are shared between the two induced samples (Fig. 2C). Among the upregulated genes are well known downstream targets of DUX4, including LEUTX, ZSCAN4, PRAMEF1 and ZNF217 (Fig. 2E). We next used Enrichr^72,73^ to search for studies that have detected similar transcriptome changes as those found in our DUX4-induction experiment, and found one other DUX4 study that has been entered into the Enrichr database (GSE33799)^10^ that shows high similarities to our data. Based on Enrichr entries, the upregulated genes in induced DIE cells are linked to DUX4 activity [-Log10(P-value) = 100.3], as are the downregulated genes [-Log10(P-value) = 3.8] (Fig. 2F and table S2). Interestingly, PAX5 appears to also be linked to the upregulation of some of the same genes as DUX4 (21 and 24 overlapping genes). The Paired box (PAX) family of genes has previously been linked to FSHD and DUX4, and could explain this overlap in gene activation^17,74,75^. Additionally, genes are often regulated by more than 1 transcription factor, which could also explain the shared target genes. We next compared the list of differentially expressed genes (DIE_8.5h) with 4 other studies that have previously explored the DUX4 transcriptional network (Geng, Rickard, Jagannathan and Heuvel)^10,20–22^. As shown in Figure 2G, we observed a high percentage of overlap between our dataset and those previously reported. The overlap between the upregulated genes is greater than the overlap that can be seen in the downregulated genes. This applies not only for the overlaps seen between our data and the other datasets, but also between the 4 other datasets (Fig. 2G and table S3). This confirms the believe that DUX4 is an activating factor, and that the down regulated genes might be more influenced by cell type.

**Figure 2.**
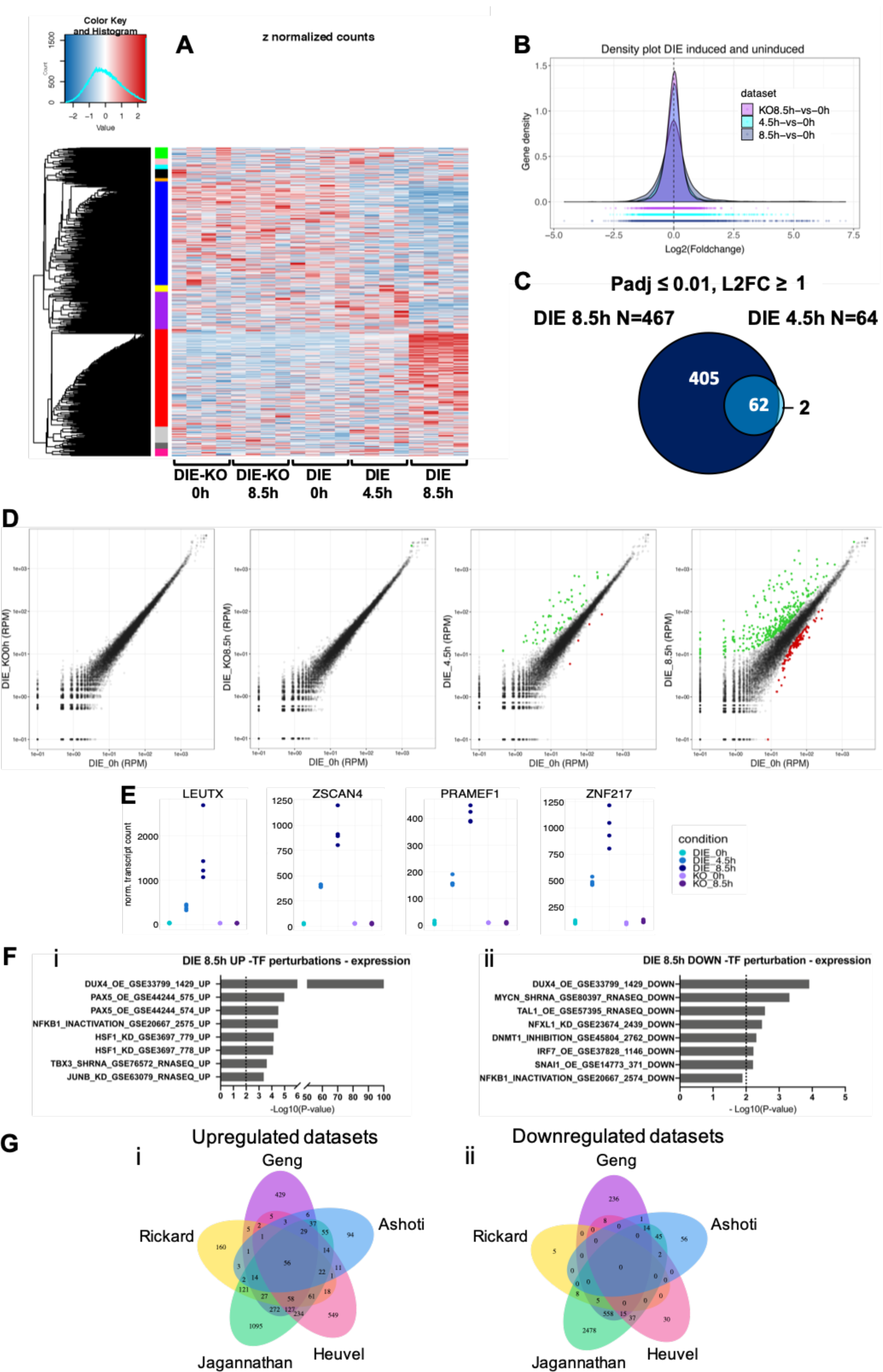
RNA-sequencing data reveals differentially expressed genes upon DUX4 expression. **(A)** Heatmap showing differentially expressed (DE) genes between samples, with gene clusters (color coded) on y-axis, and samples on the X-axis. **(B)** Gene density plot demonstrating the effects of DUX4 activation on the transcriptome of the DIE cell line. DUX4 activation results in an increase of DE genes compared to uninduced DIE cells, as indicated by the bell shape widening and shortening. **(C)** Venn diagram showing the overlap and the number of DE expressed genes at 4.5h and 8.5h of doxycycline induction (Adjusted P-value ≤ 0.01, and absolute Log2FC ≥ 1). **(D)** Scatter plots of gene expression (RPM: reads per million) of induced DIE cells versus uninduced DIE cells. Left two panels demonstrate uninduced DIE cells (DIE_0h) on the x-axis versus uninduced or induced DIE-KO samples (KO0h and KO8.5h) on the y-axis. Right two panels compare the uninduced DIE cells on the x-axis with induced DIE samples (4.5h and 8.5h) on the y-axis. Green and red points represent the DE genes with an Adjusted P-value ≤ 0.01, and absolute Log2FC ≥ 1. Green points represent upregulated genes, and the red points represent downregulated genes. **(E)** Count plots showing UMI and between sample normalized transcript counts of 4 known DUX4 targets genes: LEUTX, ZSCAN4, PRAMEF1 and ZNF217, in uninduced and induced DIE and DIE-KO cells. Every sample contains 4 points, representative of the 4 technical replicates. **(F)** TF perturbations analysis identifying transcription factors that are linked to the i) upregulation and ii) downregulation of the DE genes found in this study, by the activation or inhibition of such TFs. Activation: OE or ACTIVATION, inhibition: KO, KD, SIRNA, SHRNA, INACTIVATION, or INHIBITION. **(G)** Quintuple Venn diagram comparing DUX4 i) upregulated and ii) downregulated genes found in this study (Ashoti) to those found in previous transcriptomic studies (Geng with P-value ≤ 0.01, FDR ≤ 0.05, abs L2FC ≥ 1; Rickard with Padj value of < 0.005 and abs L2FC > 2; Jagannathan with P-value ≤ 0.01, FDR ≤ 0.05, abs L2FC ≥ 1, Heuvel with P-value ≤ 0.005, FDR ≤ 0.05, abs L2FC ≥ 1).

Overlapping data seen here are likely an underrepresentation, due to the presence of gene families containing paralogs and pseudogenes in either reference genome databases, which can lead to multi-mapped or ambiguous reads^76^. To conclude, data shown here strongly suggest that in our DIE cell system, DUX4 induces transcriptional changes similar to those found in FSHD muscle.

### Genome-wide CRISPR Screen reveals large chromosomal truncations

Using our DIE cell system, we sought out to identify modulators of DUX4 cytotoxicity by performing a genome wide CRISPR/Cas9 knockout screen. The Brunello human CRISPR knockout pooled library was used for this purpose^55^. This library contains 77.441 gRNAs targeting all protein coding genes, with an average of 4 gRNAs per gene as well as 1000 non-coding control gRNAs. To optimize the signal-to-noise ratio of the experimental system, we titrated the timing and dose of the doxycycline-mediated DUX4 induction and selected two conditions, low (250ng/ml) and high (1000ng/ml) doxycycline with an exposure time of 24h (Fig. 3A). At these concentrations 95 to 99 % of the cells die respectively. Figure 3B outlines the setup of the screen. In addition to the high and low doxycycline concentrations, cells were harvested at two timepoints after doxycycline exposure to allow recovery, early (24h) and late (72h), ultimately resulting in 4 separate 4 screens; low doxycycline/early harvest, low doxycycline/late harvest, high doxycycline/early harvest, and high doxycycline/late harvest.

**Figure 3.**
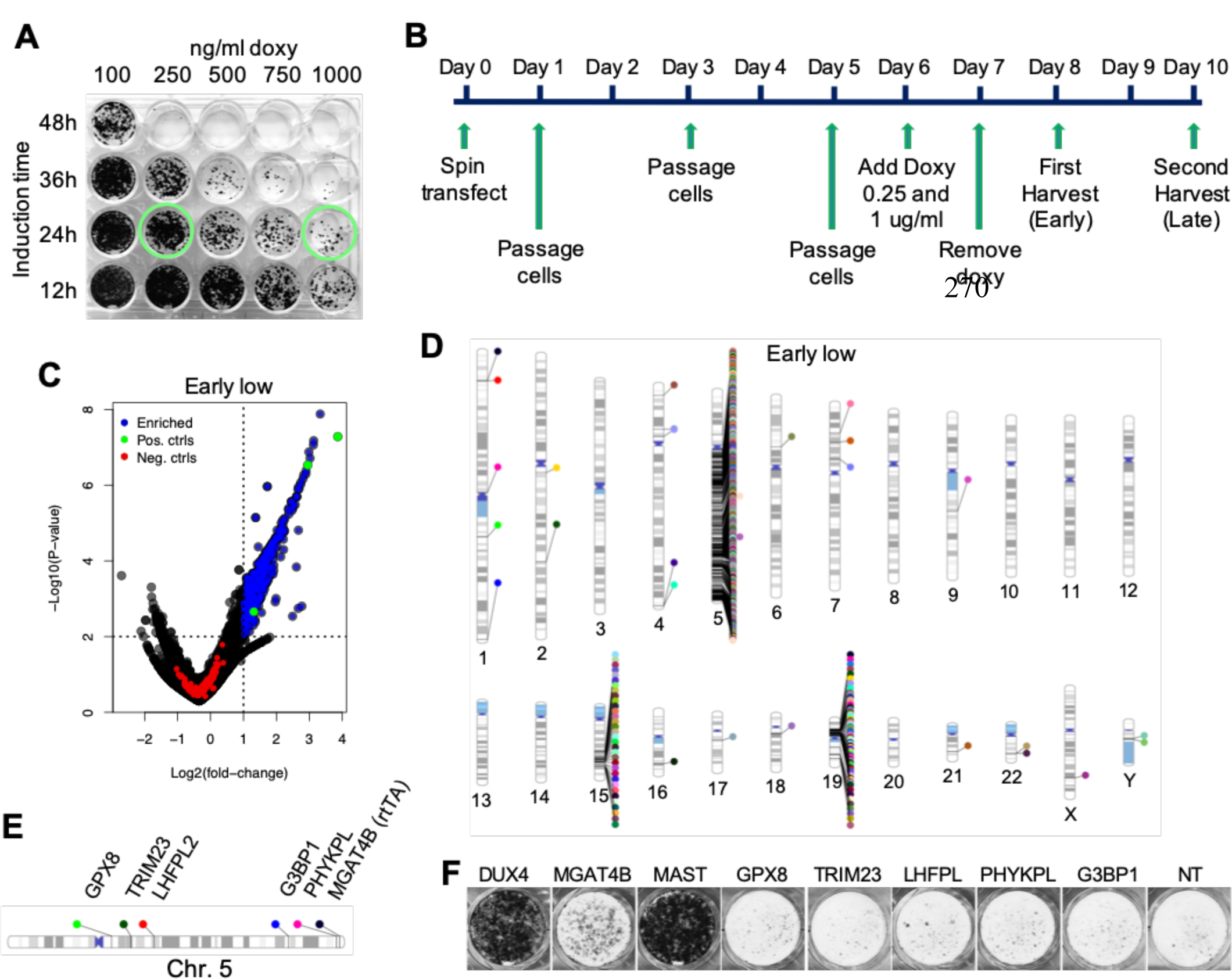
CRISPR Screen set up and discovery of a Cas9 artefact. **(A)** Viability staining of DIE cells treated with doxycycline titration curve to determine which concentration and exposure time to use to induce sufficient cell death rates in DIE cells. Green circles indicate which conditions were used for the genome wide CRIPSR/Cas9 screen. **(B)** The CRISPR/Cas9 screen timeline from the moment of library transfection (Day 0) to the final harvest of surviving DIE cells (Day 10). **(C)** Volcano plot showing the enrichment of sets of guides of the low doxycycline-early harvest screen. For a two-sided analysis see supplementary figure S6. Data shown here shows the average log2FC and log10PV of each guide set (set: 4 guides per gene). The Log2(foldchange) is plotted on the X-axis and the -Log10(p-value) on the Y-axis. Blue points represent guide sets that are significantly enriched (P-value ≤ 0.01), LFC ≥ 1), green point are the positive controls (DUX4, MAST1, MGAT4B), red points represent the Non-Target/negative control guides. **(D)** Chromosomal ideogram indicating the location of enriched hits in the human genome, of the low doxycycline-early harvest screen. **(E)** Schematic representation of the location of a small number of false positive hits on chromosome 5. **(F)** Viability staining demonstrating surviving DIE-Cas9 cells containing knock-outs of the same genes mentioned in (C), after 250ng/ml doxycycline exposure. Media did not contain any selection markers. NT: Non-Target controls.

Upon doxycycline administration and induction of DUX4 expression, cells from the surviving populations were harvested and genomic DNA was sent for sequencing. Sequencing results of the treated samples revealed a large number of significantly enriched hits (Fig. 3C and S3 for all 4 screens), including DUX4 itself and some others performing as well as the DUX4 gRNAs. However, upon closer examination it became clear that the majority of these enriched guides were located on the q arm of chromosome 5, suggesting an FSHD unrelated, experimental artefact. Since the rtTA transgene responsible for DUX4 induction is located on the 5q arm, it is likely that when Cas9 is being targeted to the q-arm of chromosome 5 it leads to the removal of the rtTA transgene, potentially through generation of a large deletion, chromosomal truncation or chromosomal rearrangement. It appears that as the rtTA integration site is located at the end of chromosome 5q, each target upstream of this site (towards the centromere) can cause a Cas9-mediated truncation, thereby removing the rtTA. (Fig. 3D, for phenograms of all 4 screens see S4). The correlation between the significance of a hit and its position along chromosome 5 highlights the strong association of these unexpected chromosomal rearrangements and the integration of rtTA at the end of chromosome 5, where the most significant hits reside in all four screens (Supplementary Figure S5).

Some of these 5q locating guides (Fig. 3E) were tested individually in DIE cells containing a constitutively expressing Cas9 its genome (DIE-Cas9), and without selecting for the rtTA and DUX4 transgene. No increased survival was detected compared to the background surviving cells that are seen in the non-target control situation (Fig. 3F). This suggests that the Cas9-induced truncation of a chromosomal arm and subsequent removal of rtTA activity is a rare event that was only identified due to the high sensitivity of our screen.

Data shown here was analyzed one-sidedly, and only truly represent enrichment. When analyzing the screen data double sided, one can again notice a clear enrichment of gRNA sequences, however no real depletion (Fig. S6).

### Filtered Genome wide CRISPR screen results reveal no single targetable gene

Since potential hits were likely obscured by the large number of false-positive hits that resulted from Cas9-mediated elimination of either the DUX4 or the rtTA transgenes, we filtered the screen results to remove all hits located on the q-arm of chromosome 5, or the p-arm of chromosome 19 (Figure 4A). After analyzing individual guides for their apparent effectiveness in the genome wide screen (instead of the group average), a list of potential hits emerged (P ≤ 0.05, Log2foldchange ≥ 1) for each of the 4 screens. Figure 4B shows the number of potential hits that met these criteria for each screen and how many of these hits are shared between them (See Table S4 for the full lists of hits). We further focused on hits that emerged in at least 3 out of the 4 screens. Hits were validated by performing individual knock outs in the DIE-Cas9 cells, now also containing an inducible eGFP in its genome (DIE-Cas9-ieGFP). The TRE controlling eGFP expression is identical to the TRE controlling DUX4 expression. If there is a true target that can mitigate the apoptotic phenotype without interfering with the inducible system, these positively targeted cells should not only survive but also emit an eGFP signal upon doxycycline admission (Fig. S7). Results show that MED25 increased cell survival when knocked-out (Fig. 4C). MED25 is a subunit of Mediator, a large complex that functions as a bridge between transcription factors and the transcriptional machinery. This includes RNA polymerase II, needed for the transcription of all protein coding genes in eukaryotes (reviewed by Soutourina^77^). The rescue seen in doxycycline induced DIE cells after MED25 knock-out diminishes upon higher doxycycline exposure, suggesting that loss of MED25 provides a partial rescue. Other genes belonging to the same mediator complex, that initially didn’t meet our criteria, were reevaluated by lowering the parameters (P ≤ 0.05, foldchange of ≥ 1.5), identifying a number of other subunits of the mediator complex. When individual knock-outs of these genes were performed, two more subunits of the Mediator-complex showed partial rescue (Fig. 4D). Finally, the individually tested KO cells were analyzed by flow cytometry, for the detection of eGFP. FACS analysis reveals that Mediator-complex components have a general effect on the inducible transcription of DUX4, since the knock-out of Mediator genes also reduced rtTA-inducible eGFP expression, suggesting that their survival was due to a generally reduced ability of rtTA to mediated transgene activation (Fig. 4E).

**Figure 4.**
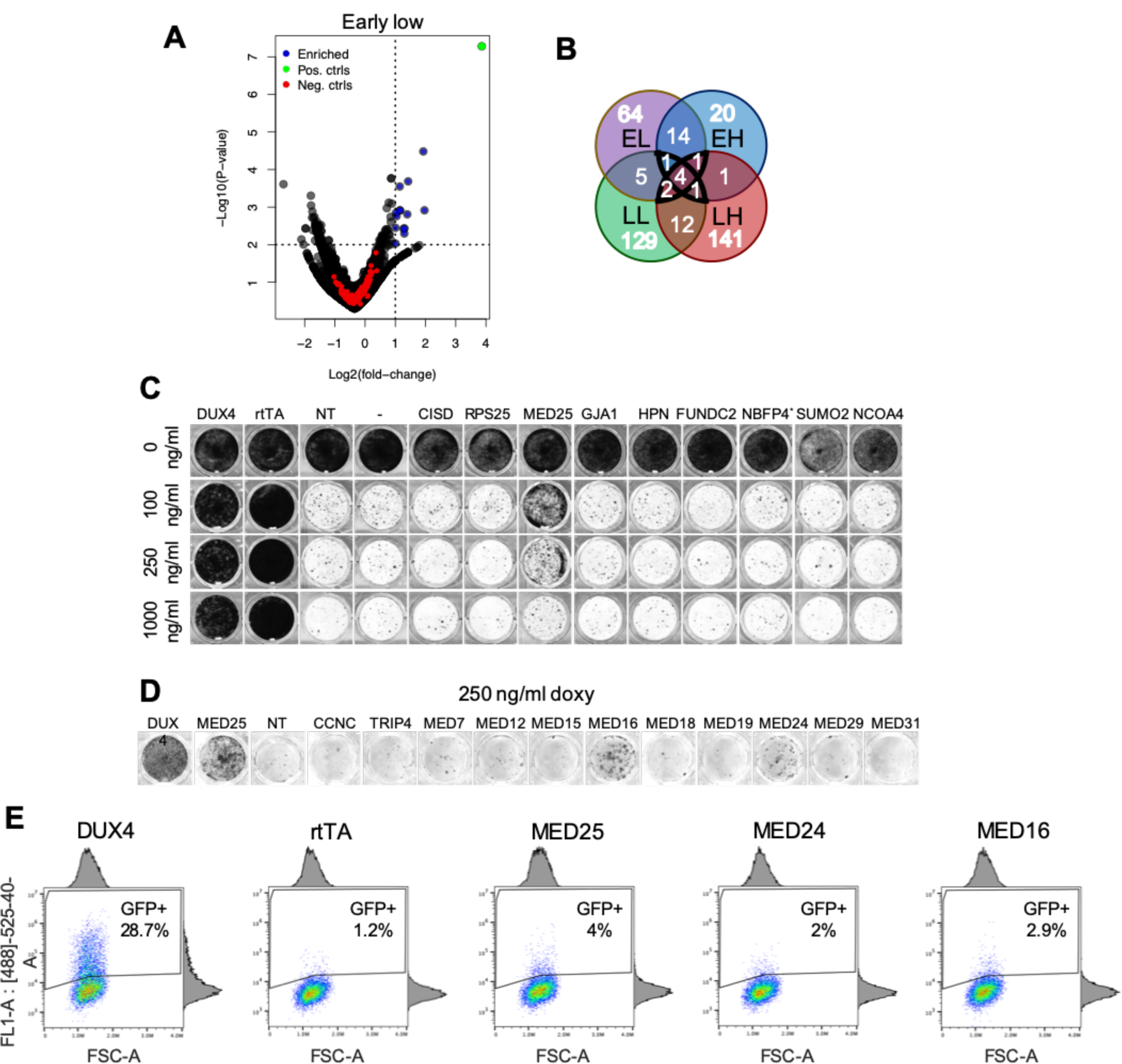
Filtered CRISPR screen data and validation of potential hits. **(A)** Adjusted volcano plot of low doxycycline-early harvest screen showing the enrichment of sets of guides targeting genes not located on chromosome 5q or chromosome 19p. Blue points represent guide sets that are significantly enriched (P-value ≤ 0.01), LFC ≥ 1), the green point is the positive control (DUX4), red points represent the Non-Target control guides. **(B)** Venn diagram showing the overlap of filtered hits between the four screens (EL: Early harvest-Low doxy, LL: Late harvest-Low doxy, EH: Early harvest-High doxy, LH: Late harvest-High doxy). **(C)** Viability staining showing surviving DIE cells containing single knockouts of potentials hits, identified in the CRISPR screen, after exposure to 3 different concentrations of doxycycline. **(D)** Viability staining showing the surviving DIE-ieGFP-Cas9 cells containing single knockouts of mediator complex subunits, after exposure to 250ng/ml doxycycline. **(E)** FACs data showing GFP positive cells in surviving populations of DIE-ieGFP-Cas9 cells containing single knock-outs. DUX4 knock outed DIE-ieGFP-Cas9 cells comprise 42% of eGFP positive cells. rtTA, MED25, MED24 and MED16 knock-outs show little eGFP expressing cells, containing between 1.2-4% of eGFP expressing cells.

In a recent study by Shadle and colleagues, a siRNA screen was performed targeting the “druggable” genome to identify pathways of DUX4 toxicity. The study revealed the MYC-mediated apoptotic pathway and the viral dsRNA-mediated innate immune response pathway to be involved in DUX4 induced apoptosis^78^. We examined our data for enrichment of gRNA sequences that target the genes identified in the Shadle study, but did not observe significant enrichment in our CRIPSR screen data of these sequences. Figure 5A shows data plots that display the enrichment (L2FC) and significance (-Log10(P-value)) of DUX4 and 3 other genes that were initially considered hits, yet later single knock-outs validations demonstrated them to either have a generally effect on transcription (MED25), or upon their knockout not exhibiting any additional survival in induced DIE cells (RPS25 and CISD). Genes were only considered if a minimum of one gRNA showed significant enrichment in at least 3 out of 4 screens. Genes involved in the pathways identified by Shadle et all. did not meet these criteria (Fig. 5B). Furthermore, knocking out these genes in the DIE cells did not show an increased survival compared to background noise (Fig. 5C, top panel), as is noticeable in some of the false positives identified during this CRIPSR screen (Fig. 5C, lower panel). It should be mentioned that the two screens have major technical differences, such as screening method, complete or partial loss of function, different scale and different cellular backgrounds, most likely all attributing to the little correlation seen between the two studies.

**Figure 5.**
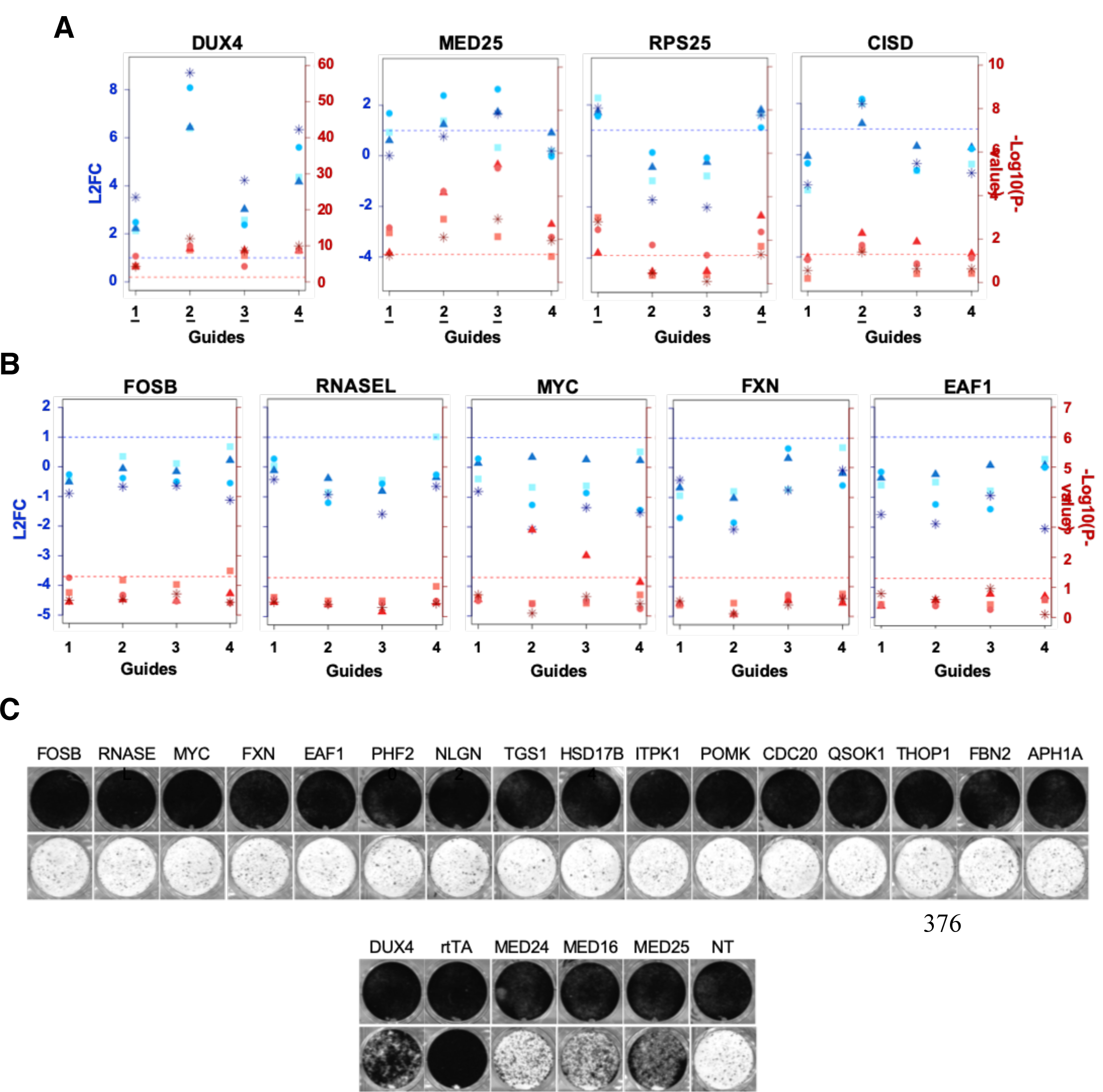
Validation of genes involved in the MYC-mediated apoptotic pathway and the viral dsRNA-mediated innate immune response pathway. **(A)** Data plots showing the significance and enrichment of gRNAs targeting DUX4, MED25, RPS25 and CISD. The Log2(fold-change) of each individual guide is plotted on the left y-axis indicated in blue, and the –Log10(P-value) is plotted on the right y-axis, in red. When guides fall above the blue and red intermitted ablines, they are considered significant (Log2(foldchange) > 1, -Log10(P-value) > 1.3). The gRNAs that are significantly enriched in all 4 screens are underlined. All 4 gRNAs targeting DUX4 are significantly enriched. 3 out of 4 gRNA’s targeting MED25 are significantly enriched (guides 1, 2 and 3). Guides 1 and 4 targeting PRS25 are significantly enriched, and CISD has one guide that is significantly enriched in all 4 screens. **(B)** Data plots showing the enrichment of gRNA targeting FOSB, RNASEL, MYC, FXN and EAF1. None of the 4 guides show significant enrichment in any of the 4 screens. **(C)** Viability staining showing surviving DIE-Cas9 cells containing single knockouts of genes involved in the MYC-mediated apoptotic pathway and the dsRNA-mediated immune response pathway (Top panel). Controls can be found in the bottom panel and are as followed, positive controls: DUX4, rtTA, MED24, MED16 and MED25; Negative non-target control: NT.

## Discussion

At present there are no effective pharmacological treatment options that can improve muscle strength or slow down disease progression in FSHD patients^79^. Unravelling the underlying mechanism of DUX4 cytotoxicity would help identify therapeutic targets. We hypothesized that inhibition of key downstream DUX4 effectors would slow or abrogate the cytotoxic process, and set out to identify such genes by performing an exhaustive genome wide CRISPR/Cas9 screen. We know the screen was exhaustive because we picked up rare rearrangements disabling the DUX4 transgene or the rtTA inducer. The goal of the screen was to identify targets that can mitigate DUX4 induced toxicity. While the screen’s technical execution went very well and displayed high sensitivity and specificity toward candidate editing events that indeed mitigated cytotoxicity in our transgene model, none of the obtained hits had a direct effect on DUX4 or its downstream transcriptional network. Rather, screen hits seemed to specifically affect the experimental system itself, either by affecting the tetracycline-inducible system responsible for DUX4 transgene induction, or by mutations of the DUX4 transgene. The main contributor of the false positive result is likely a rare Cas9-induced chromosomal truncation event, that removes the transgenes when targeted to the chromosomal arm to which they have integrated. Although these events appear to be rare, nearly all guides that targeted genes located on the chromosomal arm to which rtTA had integrated (5q) were robustly enriched, underwriting the sensitivity of this screening method. Most remaining hits did not appear to effect DIE cell survival upon individual validation, but members of the Mediator complex did show a positive effect on survival. Unfortunately, these mediator subunit genes seemed to generally suppress rtTA-mediated transcription so their mitigating effect was not mediated by specifically altering DUX4 cytotoxicity. We therefore concluded, that based on the conditions used in this study, there to be no individual target (other than DUX4 itself) that upon knockout, can provide a strong inhibition of DUX4 induced cytotoxicity. We therefore believ that efforts should therefore be redirected to the direct modulation of DUX4.

While our library only targeted protein-coding genes, we believe we would have picked-up any mitigating non-coding RNAs as well, had they provided a strong rescue from the DUX4 cytotoxic effects. In that case, one would have expected to see a similar hotspot of gRNAs on and around the true target sites, as we observed for, MED16, where a hotspot of gRNAs was observed on the p-arm of chromosome 19, corresponding to the location of MED16. Another hotspot can be seen on the q arm of chromosome 19, corresponding to the location of MED25. The hotspot on chromosome 15 can be explained by the genetic makeup of the HAP1 cell line. HAP1 cells not only have the Philadelphia chromosome, but also an integration event where a region of chromosome 15 integrated on chromosome 19p^80^. The hotspot on chromosome 15 correlates to the region that has integrated on chromosome 19p, close to the MED25 site.

Our screen results shown here do not corroborate previous findings of Shadle et al^78^. However, their siRNA screen differs in many aspects to the performed genome wide CRIPSR/Cas9 screen we executed. Their screen was knocking-down the druggable genome, using Lipofectamine RNAiMAX to deliver the siRNA library, in Rhabdomyosarcoma derived cells; whereas our screen was knocking-out protein coding genes genome wide, using a viral library, in chronic myeloid leukemia derived cells. These differences could explain why results between the two screens are not correlating with one another. Furthermore, A side by side comparison study of CRISPR/Cas9 and a next generation RNAi screens reveals that the screening methods seem to effect different biological aspect of the cells, therefore finding little correlations between results. The authors also in part attribute these differences to the technical differences between the two techniques^37^.

Another recently published genome wide CRISPR/Cas9 study, where a similar methodology was used in a DUX4 inducible, immortalized myoblast line, identified the HIF1 oxidative stress pathway as a modulator of DUX4-induced apoptosis^81^. This data, as well as previous reports clearly demonstrate the role of the HIF1 hypoxia pathway in DUX4-mediated cytotoxicity^81–83^. The HIF1 pathways did not come up in our screen (Fig. S8). This demonstrates that changes in this pathway are likely not the only DUX4 induced cellular changes that push cells towards apoptosis. The fact that the HIF1 pathway did not come up in our screen could also indicate differences in sensitivity to oxidative stress between cellular systems. Different cell types experience and respond differently to oxidative stress, with differences in culturing conditions as a further attributing factor, such as the concentration of 2-mercapto ethanol to cell culture media.

While our screen did not identify target genes that can mitigate DUX4 cytotoxicity, it does illustrate some important aspects that need to be considered when performing phenotypic CRISPR/Cas9 screens. One being the likely large chromosomal truncations that can be induced Cas9, a phenomena also recently reported by Cullot et al^84^. While these are rare events in a cell population, our results demonstrate that in a sufficiently sensitive screening system, they are robustly identified and can crowd potential positive hits. Sufficient selection should at least help in this aspect by removing cells that had their resistance marker (linked to the transgene) deleted. Another aspect that needs consideration are the endogenous genes that have a general effect on transcription and translation, in this case effecting the inducible system, like subunits of the mediator complex identified in this study. Potential hits will always need to be validated individually in such a way that can exclude this possibility, like shown here, or by Shadle et al. where some of the same genes were identified effecting their inducible Tet-On system^78^.

This study started out with the aim of trying to contribute to the understanding of the underlying molecular mechanism of FSHD, by performing a genome wide CRISPR-Cas9 phenotypic screen. However, with no significant hits that can explain their contribution to the apoptotic phenotype, this story also tells a cautionary tale for knockout screens through the use CRISPR-Cas9, which can benefit future groups planning to execute similar screens.

## Methods

### Cloning and generating the DIE cell line

To generate the inducible DsRed/DUX4 system, the third generation lenti-viral plasmid pRRLsincPPT-wpre^85^ was used as the backbone. The linearized viral backbone was created by restriction digestion using the following enzymes: HpaI and SalI (NEB). All inserts were generated with PCR amplification using phusion DNA polymerase (Fischer Scientific). Insert were created with 15bp adapter sequences, matching the backbone or neighboring fragments, for in-fusion cloning (Clontech). The first fragment consisted of cPPT/CTS-TRE-mCMV sequences, and the second fragment contained the LoxP-DsRed-LoxP (LSL) sequence. After inserting these two fragments into the pRRLsincPPT-wpre backbone, this newly cloned construct was transformed into chemically competent Stbl3 Escherichia coli (E.coli). The plasmid was isolated and purified from the Stbl3 cells using the HiPure plasmid kits from Invitrogen (Fischer scientific). This TRE-LSL plasmid was then opened up using restriction enzymes XbaI and EcroI (NEB). After which the remaining three inserts: DUX4(exon1-3), mPGK and PuroR-WPRE, were cloned downstream from the LoxP-DsRed-LoxP in similar fashion.

The DIE cell line was obtained by transducing diploid HAP1 (DIPH) cells with lentiviral particles containing the inducible DsRed/DUX4 cassette mentioned above. 2 days after lentiviral transduction, positively transfected cells were selected with puromycin. After establishing a stable line by puromycin selection, lentiviral particles containing CMV-rtTA3-BlastR were added to the DsRed/DUX4 containing DIPH cells. Positively transfected cells were subsequently selected with basticidin, and FACs sorted for DsRed expression upon exposure to doxycycline. The pLenti CMV rtTA3 Blast (w756-1) was a gift from Eric Campeau (Addgene plasmid #26429).

### Cell culture

DIPH cells were cultured in IMDM media (Fischer Scientific) supplemented with 10% FBS. The DIE cells were cultured in IMDM media supplemented with 10% Tet system approved FBS (Clontech), 5μg/ml Puromycin and 6μg/ml Blasticidin, 100μM 2-mercaptoethanol.

### Cloning p2T-Cas9, p2T-ieGFP and sgRNA constructs, and generating DIE-Cas9 and DIE-Cas9-ieGFP cell lines

The p2T-CAG-spCas9-NeoR mammalian expression plasmid was created by replacing the Blasticidin resistance gene (BlastR) in the p2T-CAG-spCas9-BlastR (Addgene: 107190) ^86^ with a Neomycin resistance gene (NeoR). The p2T-CAG-spCas9-BlastR plasmid is contained in a p2Tol2 backbone^87^. The BlastR gene was removed using restriction digestion, using MfeI and AflII (NEB). Cloning the NeoR DNA fragment into the p2T-CAG-spCas9 backbone was done in similar fashion as described above. The p2T-CAG-SpCas9-BlastR was a gift from Richard Sherwood. The p2T-TetO-eGFP-HygroR plasmid was generated in a similar way as the p2T-CAG-spCas9-NeoR. In short, all sequences between transposable elements of a p2T plasmid were removed by restriction digestion using AleI and EcoRI (NEB). The TetO-eGFP-HygroR cassette was created by amplifying each subunit individually from already excising constructs, and thereafter cloned into the empty p2T backbone, using in-fusion cloning.

Both p2T-CAG-spCas9-NeoR and p2T-TetO-eGFP-HygroR were introduced in the DIE cell line by using Transposase. The p2T-CAG-spCas9-NeoR was introduced into DIE cells together with a plasmid encoding for transposase, using Polyethylenimine (PEI) transfection reagent (4ug PEI per 1ug DNA). The DIE cells were exposed to the transfection mixture for 14-16h, after which the transfection media was replaced with growth media. Geneticin g418 selection was started two days post transfection, generating the DIE-Cas9 line. The DIE-Cas9-ieGFP cell line was created by adding Transposase and p2T-TetO-eGFP-HygroR the DIE-Cas9 line, described as above. spCas9-sgRNA constructs were cloned using a plasmid containing a U6 promotor, 2 BsmBI sites with directly adjacent the tracrRNA sequence, and a Hygromycin resistance gene (made in house). This U6-2xBsmBI-Tracr-HygroR plasmid was digested with the BsmBI restriction enzymes (NEB), after which the CRISPR inserts were ligated in using T4 DNA ligase (NEB). CRISPR inserts were generated by annealing two complementary oligos containing a 4bp adapter serving as the BsmBI sticky end.

All plasmids mentioned in this study were transformed in chemically competent Stbl3 Escherichia coli (E.coli), and prepped using a HiPure plasmid Midi or Maxi kit (Invitrogen).

### RNA extraction and RT-qPCR

Cultured cells were rinsed with DPBS just prior to the additional of TRIzol reagent (Thermo Scientific). Total RNA samples were subsequently extracted by addition chloroform, and isopropanol precipitation, and finally treated with RNase free DNase I (Promega). Reverse transcription was performed using the Superscript III kit (Invitrogen) and random primers (Promega), generating cDNA. Quantitative PCR was then initiated using IQ SYBR Green Supermix (Bio-Rad 1708880), 50 ng of cDNA, and the following gene-specific primers:

- DUX4: 5’-CCCAGGTACCAGCAGACC-3’, 5’-TCCAGGAGATGTAACTCTAATCCA-3’^88^ ;
- ZSCAN4: 5’-GTGGCCACTGCAATGACAA-3’, 5’-AGCTTCCTGTCCCTGCATGT-3’^88^;
- ZNF217: 5’-AAGCCCTATGGTGGCTCC-3’, 5’-TTGATATGACACAGGCCTTTTTC-3^88^’;
- PRAMEF1: 5’-CTCCAAGGACGGTTAGTTGC-3’, 5’-AGTTCTCCAAGGGGTTCTGG-3^88^’;
- LEUTX: 5’- GGCCACGCACAAGATTTCTC-3’, 5’- TCTTGAACCAGATCTTTACTACGGA-3’;
- rtTA: 5’- CCCTGCCAATCGAGATGC-3’, 5’-CGGTATGACTTGGCGTTGTT-3’;
- eGFP: 5’-GACCACTACCAGCAGAACAC-3’, 5’- GAACTCCAGCAGGACCATG-3’^89^;
- MED25: 5’- ACCTGGGACCCTACTTCGAG-3’, 5’- ACACCACGAGGCTGTACTGG-3’.

Data was normalized to HPRT expression by using the following primer pair: 5’- CCTGGCGTCGTGATTAGTGA-3’, 5’-CGAGCAAGACGTTCAGTCCT-3’^90^.

### Protein extraction and Western blot

DIE cells were harvested by trypsinization and lysed with RIPA buffer. Total protein concentrations were determined using a Pierce BCA protein assay kit (Fischer Scientific). 20ug protein was denatured using 4x Laemmli sample buffer (Bio-rad) with 10% BME (Sigma), and boiled for 5 minutes. Samples were run on a 15% SDS-polyacrylamide gel and transferred to a PVDF membrane (Merck). Membranes were blocked for an hour using 5% BSA in TBST, and were subsequently incubated overnight with anti-DUX4 antibody [E5-5] (Abcam, ab124699) in blocking solution (5% BSA in TBST), at 4°C. Membranes were than incubated for an hour with Secondary goat anti-rabbit-HRP antibody (Santa Cruz, sc-2004), and primary rabbit mAb β-Actin HRP conjugated antibody (Cell signaling, 5125s) in blocking buffer. Chemiluminescent signal was detected using GE ImageQuant LAS 4000 imager, upon admission of Pierce ECL Plus Western Blotting substrate (Fischer Scientific).

### RNAseq sample preparation and sequencing

Cultured cells were rinsed with DPBS just prior to the additional of TRIzol reagent (Thermo Scientific). Total RNA samples were subsequently extracted by addition chloroform, and isopropanol precipitation. The library prep was performed using CEL-seq1 primers^91^ and the Life technologies Ambion kit (AM1751)^92^, and were processed using CEL-seq2 protocol^93^. Samples were sequenced using Illumina Nextseq 500, 2×75 kit, high output. Four technical replicates per samples were send for sequencing, and were sequenced to an average of 600.000 reads per replicate (combined read count of 2.4 million reads per sample). Differential expression analysis was done using the DESeq2 package^94^.

### Doxycycline titration curve

200.000 cells were seeded into wells of a 24-wells plate and incubated overnight at 5% CO_2_ and 37°C. When cells reached a density of 90-100% confluency, different concentrations of doxycycline were added to the vertical lanes (100 ng/ml, 250ng/ml, 500ng/ml, 750ng/ml, 1000 ng/ml), with the horizontal lanes experiencing different exposure times (48h, 36h, 24h, 12h). After a recovery period of 96h (after doxy exposure was ended), cells were washed with DPBS, and fixed with Methanol for 10 minutes. Giemsa stain, modified solution (Sigma) was subsequently added for 45 minutes, after which it was removed and the wells were washed with demineralized water.

### Genome-wide CRISPR screen

The screen on the DIE line was performed as previously described by Doench et al.^55^ and Sanson et al.^95^. Due to a shared selection marker between the DIE line and the all-in-one Brunello lentiviral library, transfected cells could not be selected for, thus the total number of cells was raised to 1500 cells per guide, when considering an average transfection efficiency of 30-50% in all cell lines tested by Doench et al^55^. To minimize the probability of multiple sgRNA plasmids entering one cell, we determined the transfection efficiency and calculated the MOI. With 1500 cells per guide (total of 77.441 guides), each of the three replicates contained 120*10E6 cells. These cells were spin transfected for 2h at 1000g with 82*10E6 Brunello virus particles (LentiCRISPRv2, Addgene 73179-LV, all-in-one system in which every plasmid contains SpCas9, and a guide RNA) reaching a MOI of 0.65, and a transfection efficiency of around 60% upon testing the viral library on the DIPH parental line. After transfection the 120*10E6 transfected cells (contained in 40 wells of 12-well tissue culturing plates) were trypsinized and passaged to 60 145mm TC plates. Mutagenized cells were maintained for 6 days, before inducing a set of 24 plates with either a low or high doxycycline concentration (low: 250ng/ml, high: 1000ng/ml). The remaining 12 plates were harvested for cryofreezing (7 plates) and for determining library coverage (5 plates). After a 24h doxycycline induction period, 12 plates were given a 24h recovery period (early harvest) of both the low and high doxycycline exposed sets. The remaining 24 plates received an additional 48h of recovery time (late harvest), before harvesting the surviving cells for sequencing (Fig. S1). Cell Pellets were stored at -80°C until further processing. The Human Brunello CRISPR knockout pooled lentiviral prep library was a gift from David Root and John Doench.

### Library prep, sequencing and analysis

Genomic DNA (gDNA) was isolated using NucleoSpin Blood Mini (less than 5 million cells), Midi (L) (5-20 million cells) and Maxi (XL) (more than 20 million cells) kits, depending on the size of cell pellet. Libraries were prepared and sequenced on a HiSeq2000 (Illumina) as described by Doench et al. Analysis was conducted using “STARS”, gene-ranking method to generate FDR values developed by Doench et al. that was used to generate p-values and FDR rates.^55^ Chromosomal ideogram were generated by using the PhenoGram webtool from the Ritchie lab from the university of Pennsylvania^96^.

### Individual knock outs Cell survival Giemsa staining

DIE-Cas9 and DIE-Cas9-ieGFP were seeded in a 24-well setting. Next day, when the cells had reached 70-90% confluency, cells were transfected with 500ng guide plasmid per well using 4ug PEI per 1ug DNA. During the overnight transfection no selection markers were presents in the media, however growth media was supplemented with 100U/ml pen-strep. Cells were passaged with or without selection markers during a period of 6-7, after which doxycycline was added (100, 250 or 1000 ng/ml) for a 24h period. Wells were washed with DPBS to remove dead cells and debris. Remaining cells were given the opportunity to grow out, or to perish (if they had already entered the apoptotic pathway) for an additional 48-96 hours. The wells were stained using Giemsa modified solution, as described previously.

### Flowcytometry sorting (FACS) and analysis

DIE-Cas9-ieGFP cells were induced with 250ng/ml doxycycline 24h prior to FACS analysis. After the 24h doxycycline exposure, cells were trypsinized using 0.25% Trypsin-EDTA, resuspended in iMDM media supplemented with Tet approved FBS and DAPI nuclear staining, and strained using a Cell-strainer capped tubes (Falcon). Cells were analyzed using the Beckman coulter Cytoflex S flow cytometer.

### Imaging DIE cells

Untreated and doxycyline treated DIE cells were stained with AnnexinV-Alexa Fluor 488 and PI (Thermo Scientific), by adding the staining solutions directly to growth media at a 1:50 and 1:100 ratio respectively. Cells were left to incubate overnight in growth media containing staining solution(s) and with or without doxycycline treatment. Live imaging during treatment of these cells was carried out using a Confocal microscopy (Zeiss LSM 700). Still images were taken after the overnight incubation with am EVOS Digital Color Fluorescence Miscroscope (Invitrogen).

## Data Resources

Data containing the bulk RNA sequencing samples in quadruplicate are available from the GEO data base, accession number: GSE154649.

Data containing the Genome wide CRISPR/Cas9 samples in triplicate are available from the GEO data base, accession number: GSE155034.

## Acknowledgements

This study was supported by Stichting FSHD and the SingelSwim Utrecht. The authors like to thank Nune Schelling, Peng Shang, Stefan van der Elst, en Reinier van der Linden for excellent technical assistance.

## Supplementary figures

**Supplementary figure S1.**
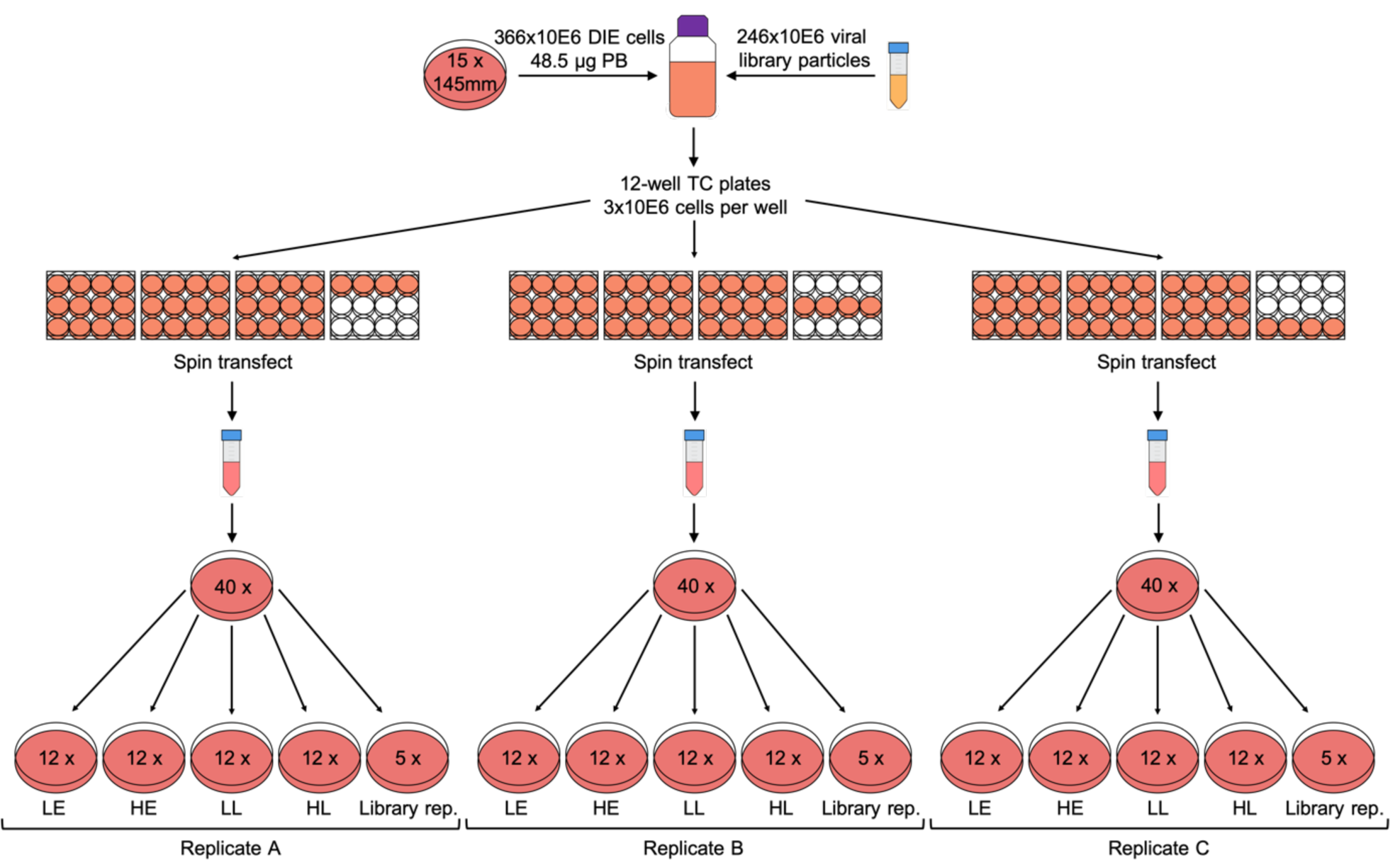
Execution of the CRIPSR/Cas9 genome wide screens. A schematic representation of the execution of the CRISPR/Cas9 screens. PB: Polybrene, TC: Tissue culture, LE: Low doxycycline/Early harvest, HE: High doxycycline/Early harvest, LL: Low doxycycline/Late harvest, HL: High doxycycline/Late harvest, Library rep: Library representation.

**Supplementary figure S2.**
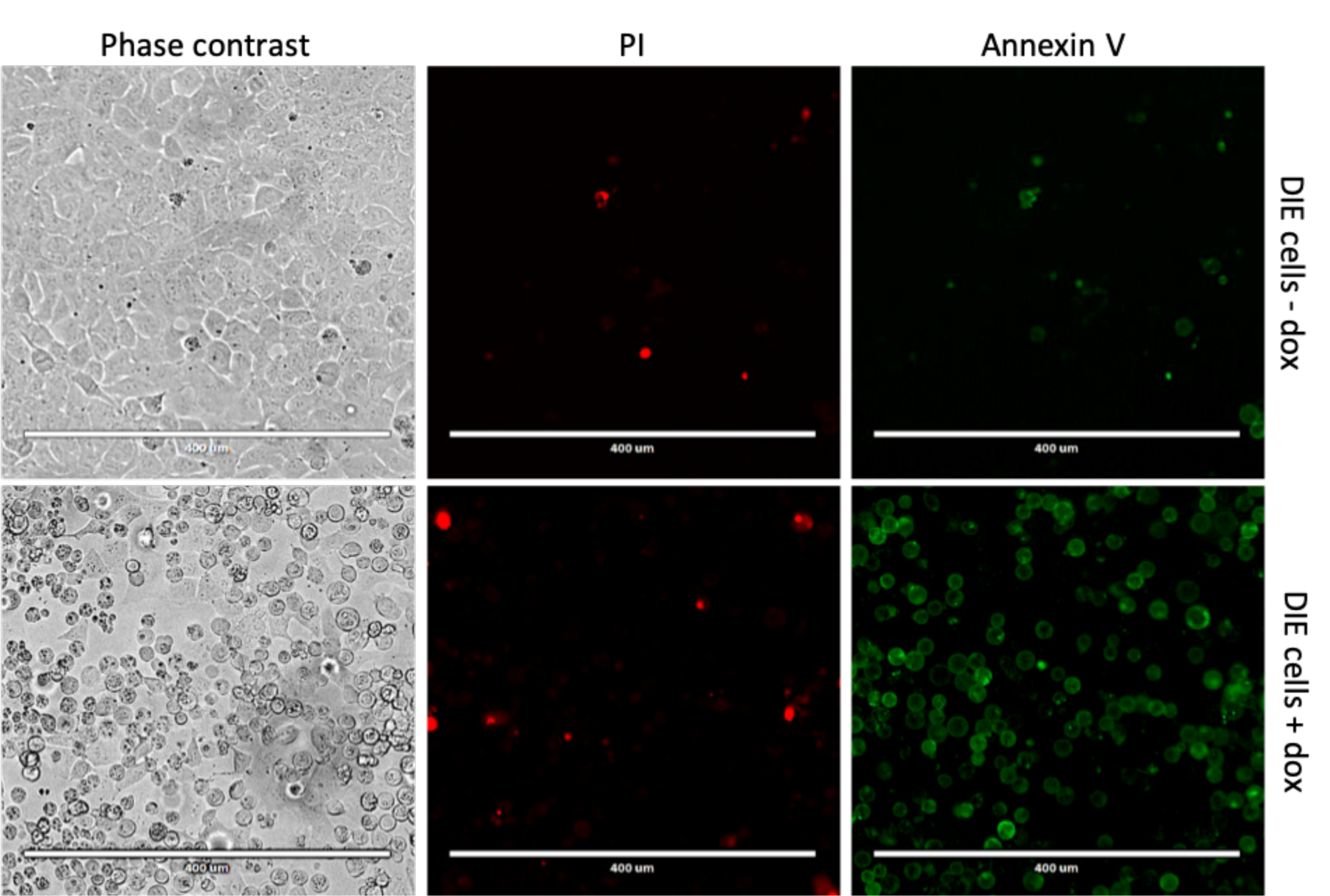
Living and dying DIE cells. Uninduced (top panel) and doxycycline induced (bottom panel) DIE cells, stained with Propidium Iodide (PI) (middle panel) and AnnexinV-Alexa Fluor 488 (right panel), with a phase contrast image in the left panel. DIE cells in the bottom panel are stained positive for AnnexinV, with no increasing PI signal compared to uninduced DIE cells (top panel).

**Supplementary figure S3.**
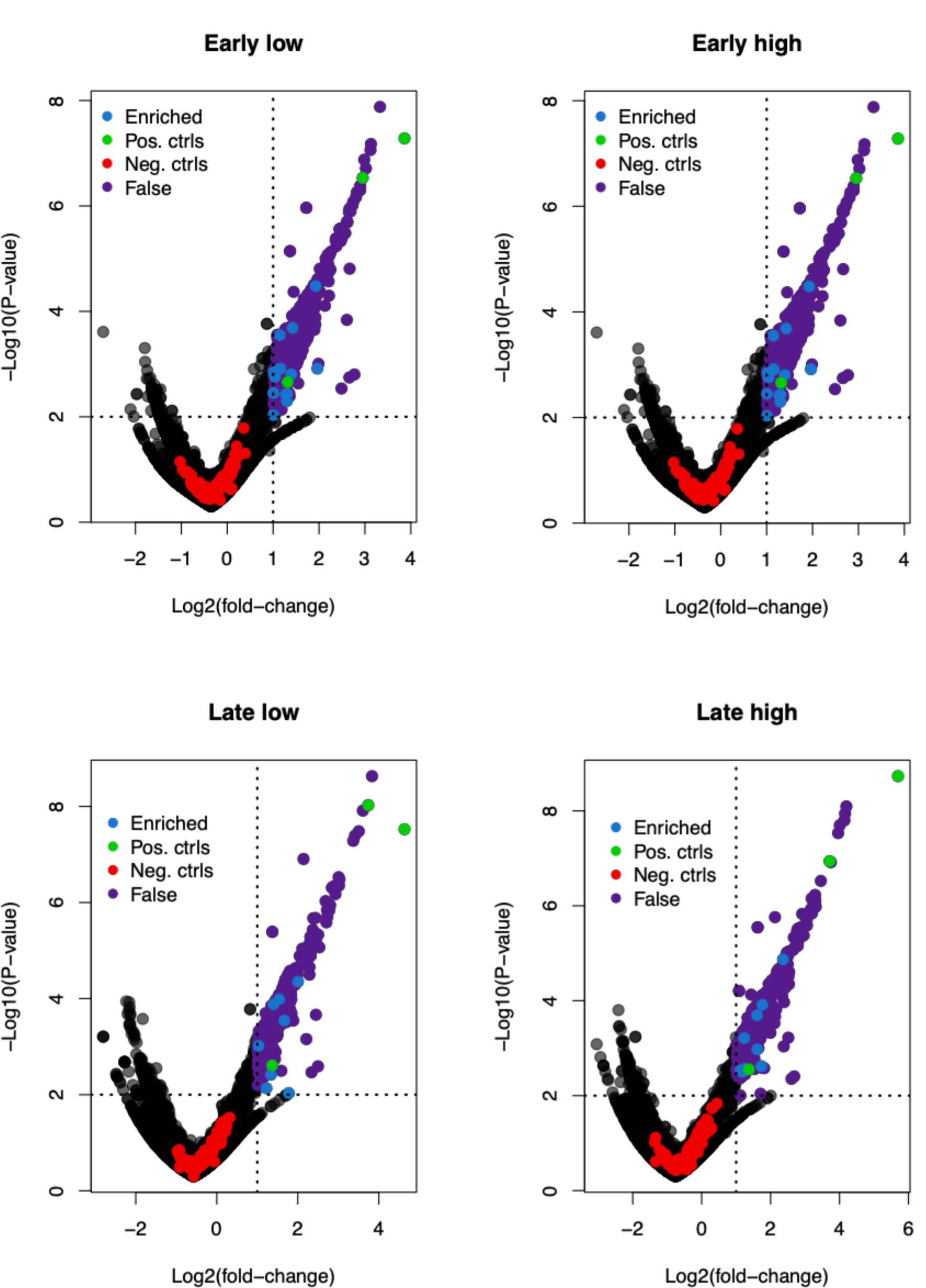
Analysis of enriched gRNAs from screen data processed with a one-sided analysis. Volcano plots illustrating enrichment of gRNAs in the surviving population of DIE cells of all 4 screens. Due to the one-sided analysis, depletion data should not be taken into consideration. For a two-sided analysis see supplementary figure S6. The Log2(foldchange) (log2FC) is plotted on the X-axis and the -Log10(p-value), (-log10PV) is plotted on the Y-axis. Data shown here shows the average log2FC and -log10PV of each guide set (set: 4 guides per gene). Blue points represent guide sets that are significantly enriched in this data set (Log2FC ≥ 1, -log10PV ≥ 2), purple points represent the false positive hits that on chromosome 5q and chromosome 19p, green point are the positive controls (DUX4, MAST1, MGAT4B), red points represent the Non-Target control guides.

**Supplementary figure S4.**
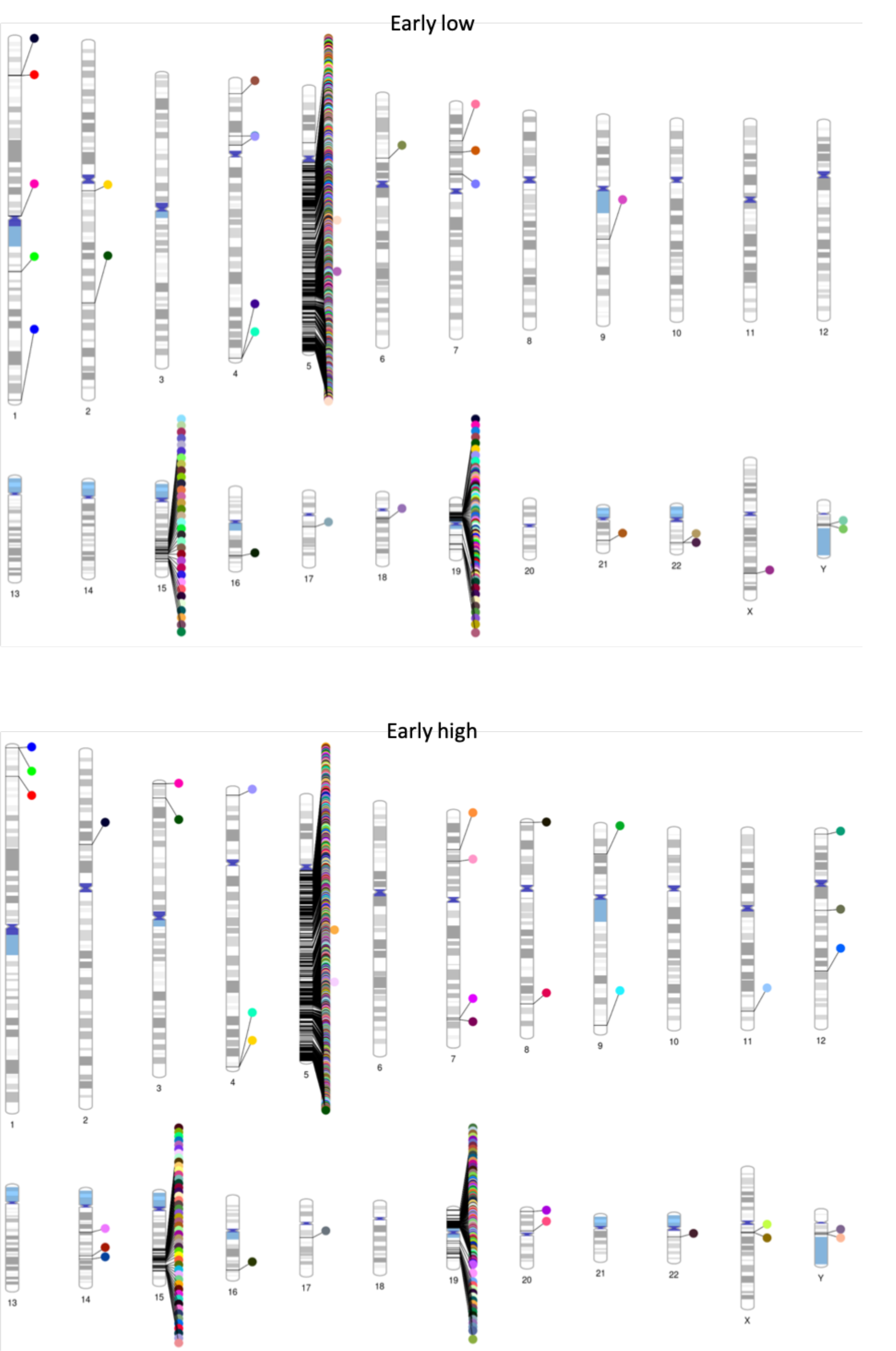
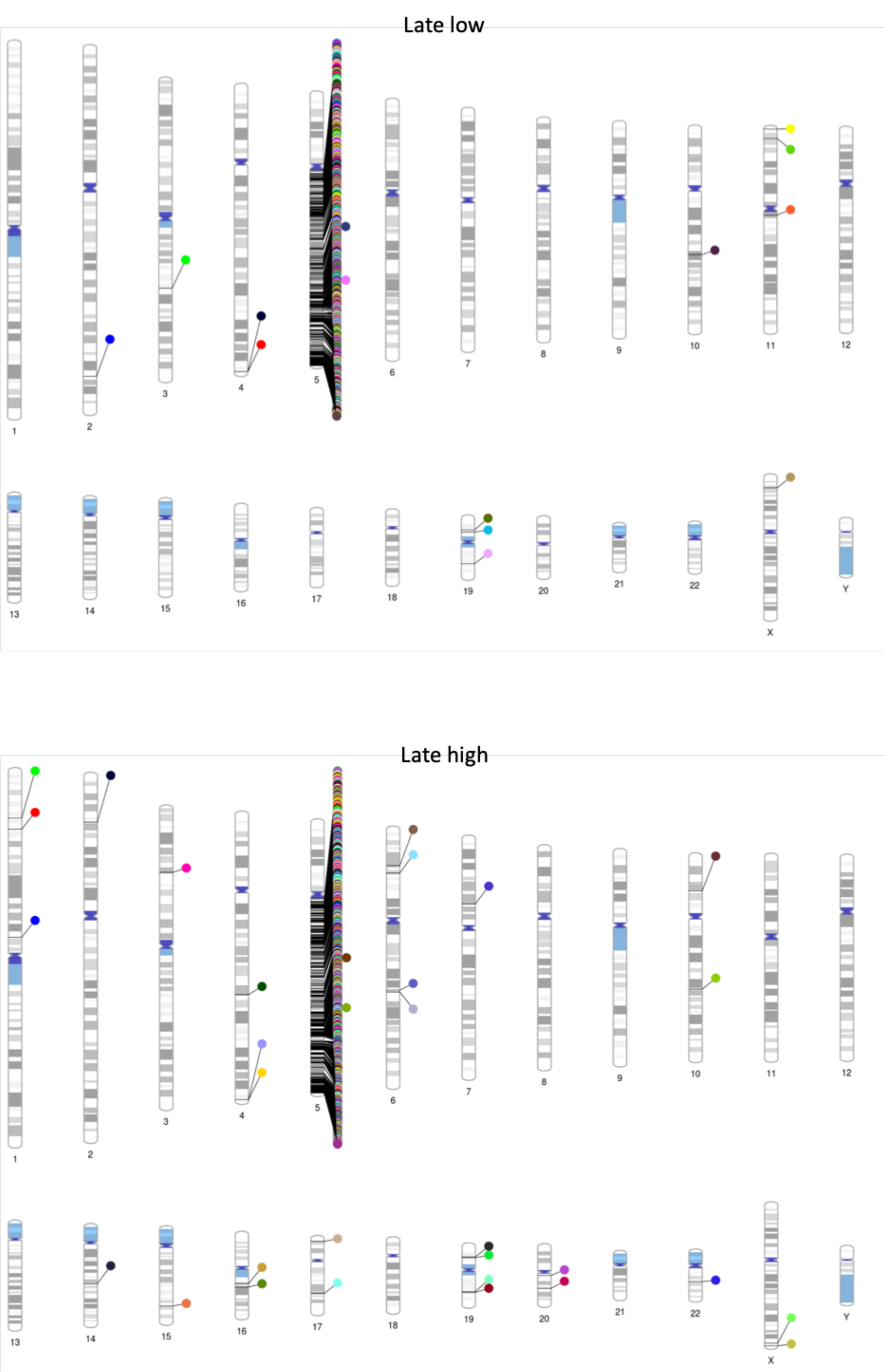
PhenoGrams showing enriched hits in the human genome. Chromosomal ideogram indicating the location of enriched hits in the human genome, for each of the 4 screens. PhenoGram is a software created by the Ritchie lab from the university of Pennsylvania^96^.

**Supplementary figure S5.**
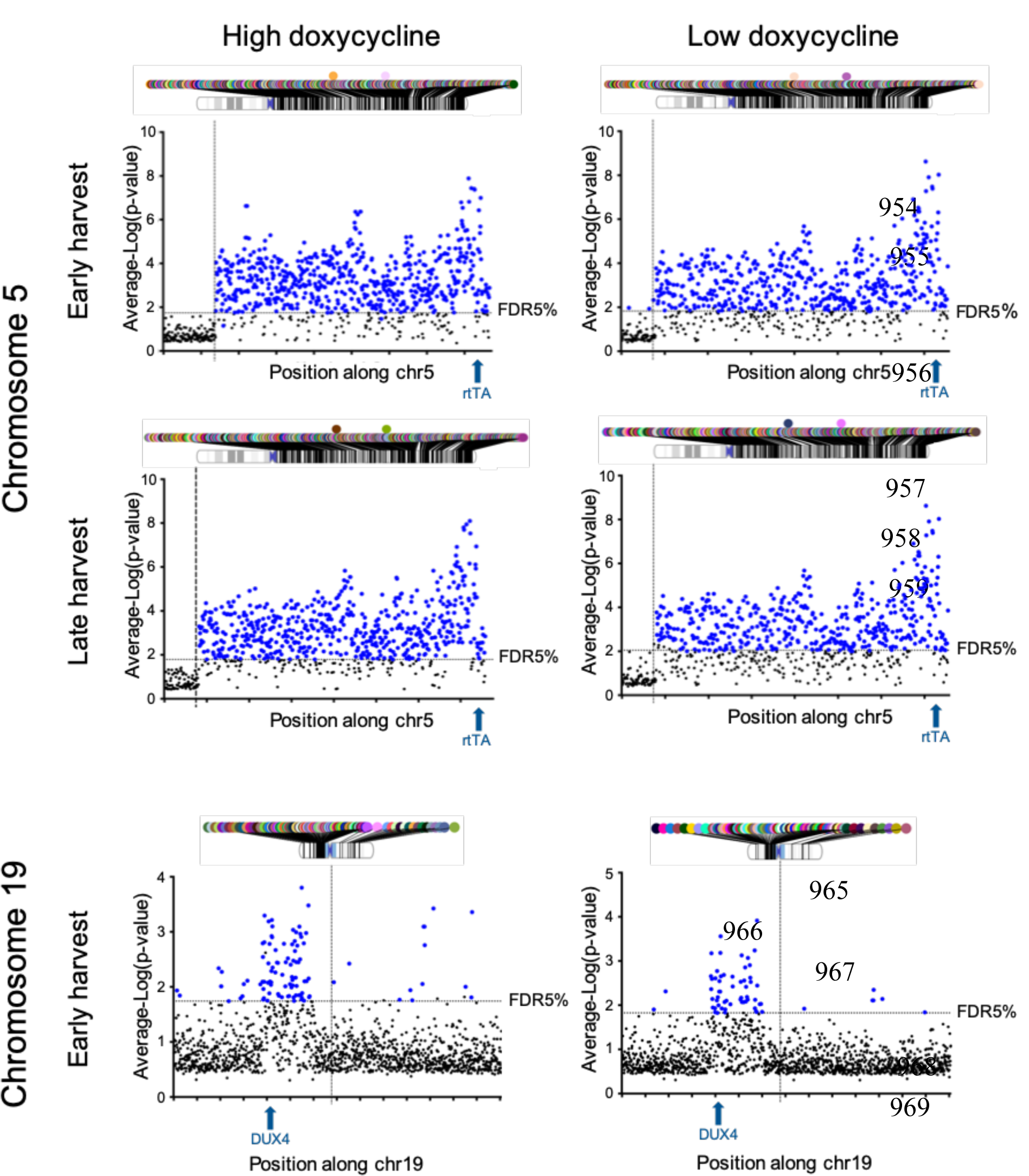
Data plot displaying enriched hits on chromosome 5q and chromosome 19. The average-Log(p-value) is plotted on the Y-axis, and the X-axis is displaying the position on the chromosome. The vertical abline indicates the position of the centromere. All points above the horizontal abline (in blue) indicating significantly enriched hits that fall below the 5% False Discovery Rate (FDR) threshold. The location of the transgene is annotated with a blue arrow on the X-axis.

**Supplementary figure S6.**
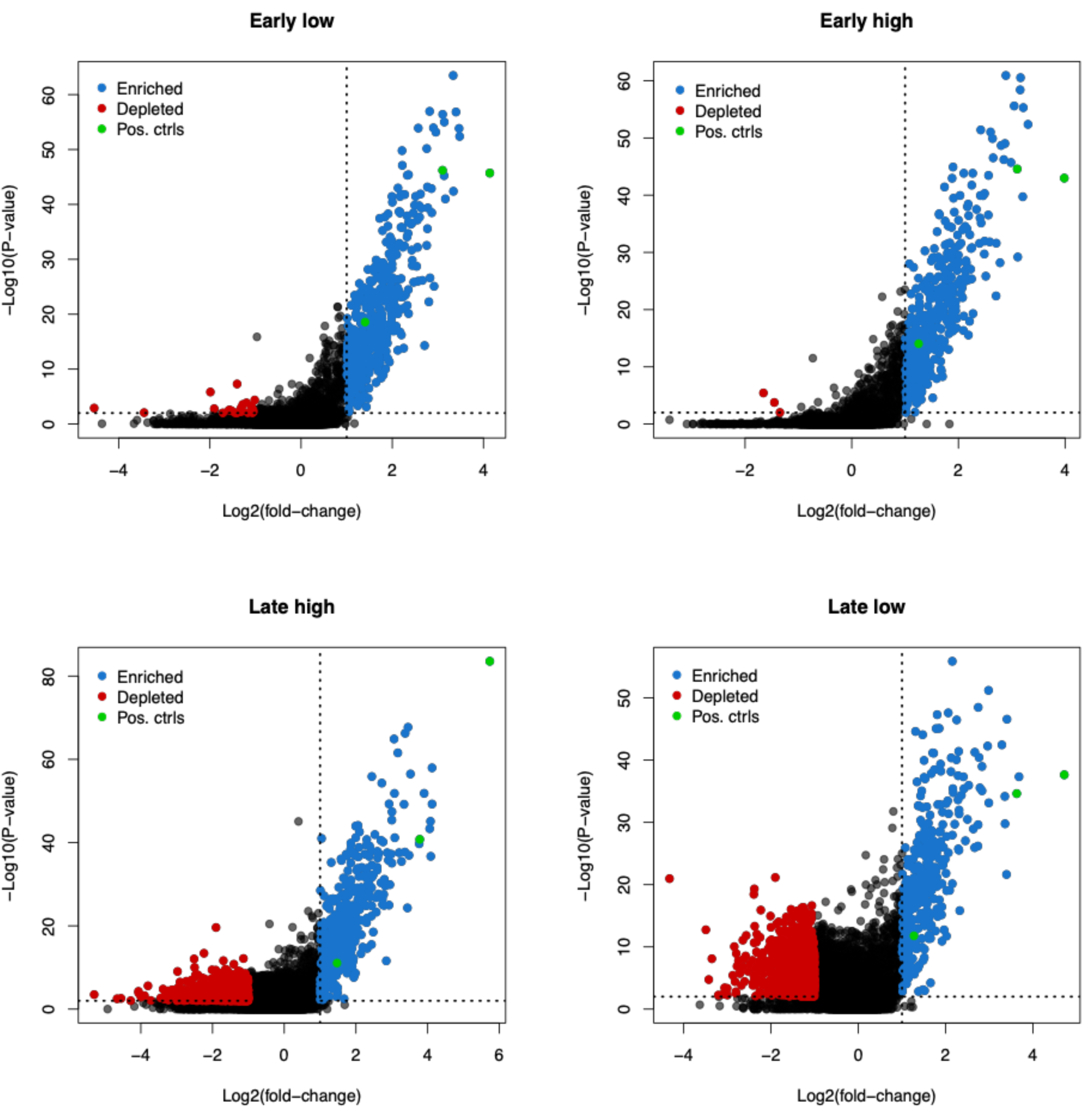
Analysis of enriched and depleted gRNAs from screen data analyzed in a two-sided analysis. Volcano plots illustrating enrichment and depletion of gRNAs in all 4 screens. The Log2(foldchange) (log2FC) is plotted on the X-axis and the -Log10(p-value), (-log10PV) is plotted on the Y-axis. Dots indicated in blue are significantly enriched gRNAs sequences (Log2FC ≥ 1, - log10PV ≥ 2). Green dots represent positive controls (DUX4, MAST1, MGAT4B), and red dots represent the depleted targets.

**Supplementary figure S7.**
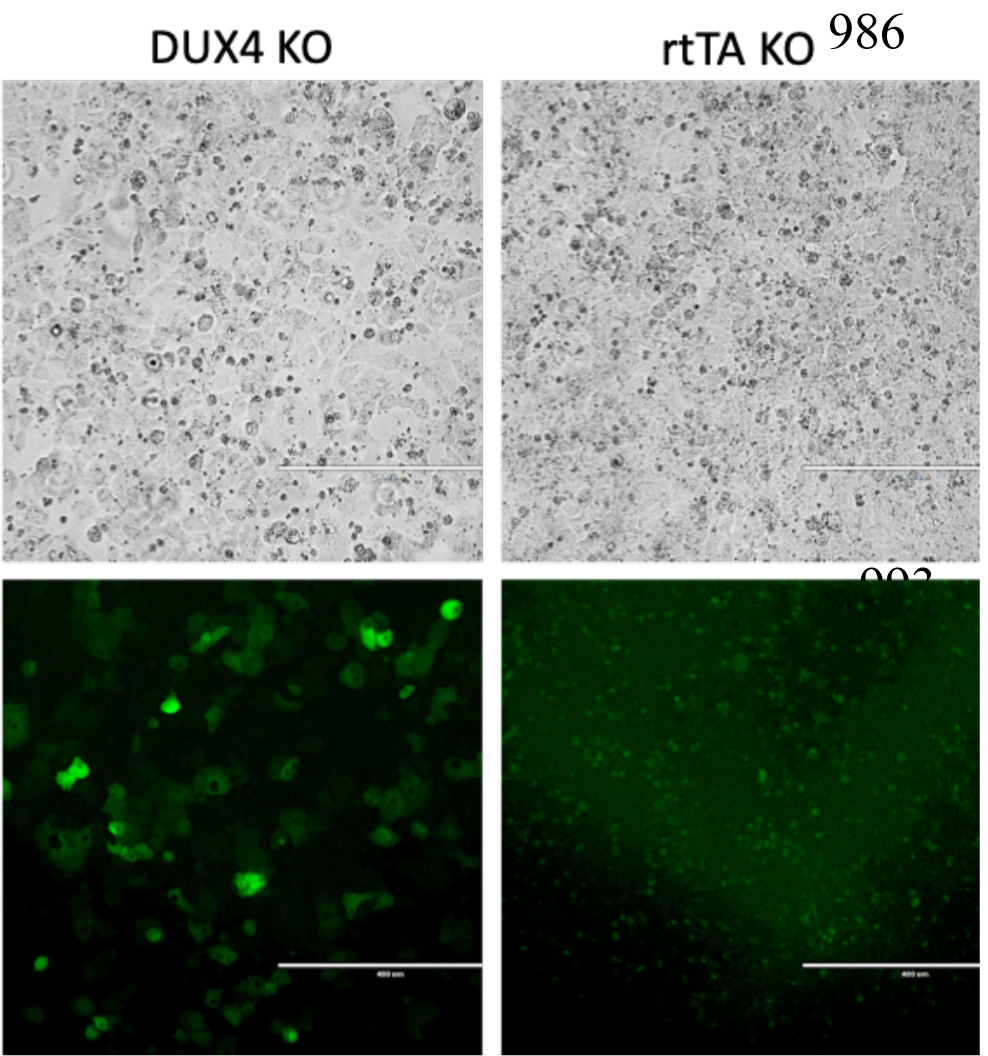
Individual knock-outs in DIE-ieGFP-Cas9 cells demonstrating eGFP activation in cells with a functional TetO inducible system. Phase contrast (top panel) and fluorescent images (bottom panel) of DIE cells containing a DUX4 KO (left panel), and rtTA KO (right panel) induced with 250 ng/ml doxycycline. DIE cells had reached over 100% confluency.

**Supplementary figure S8.**
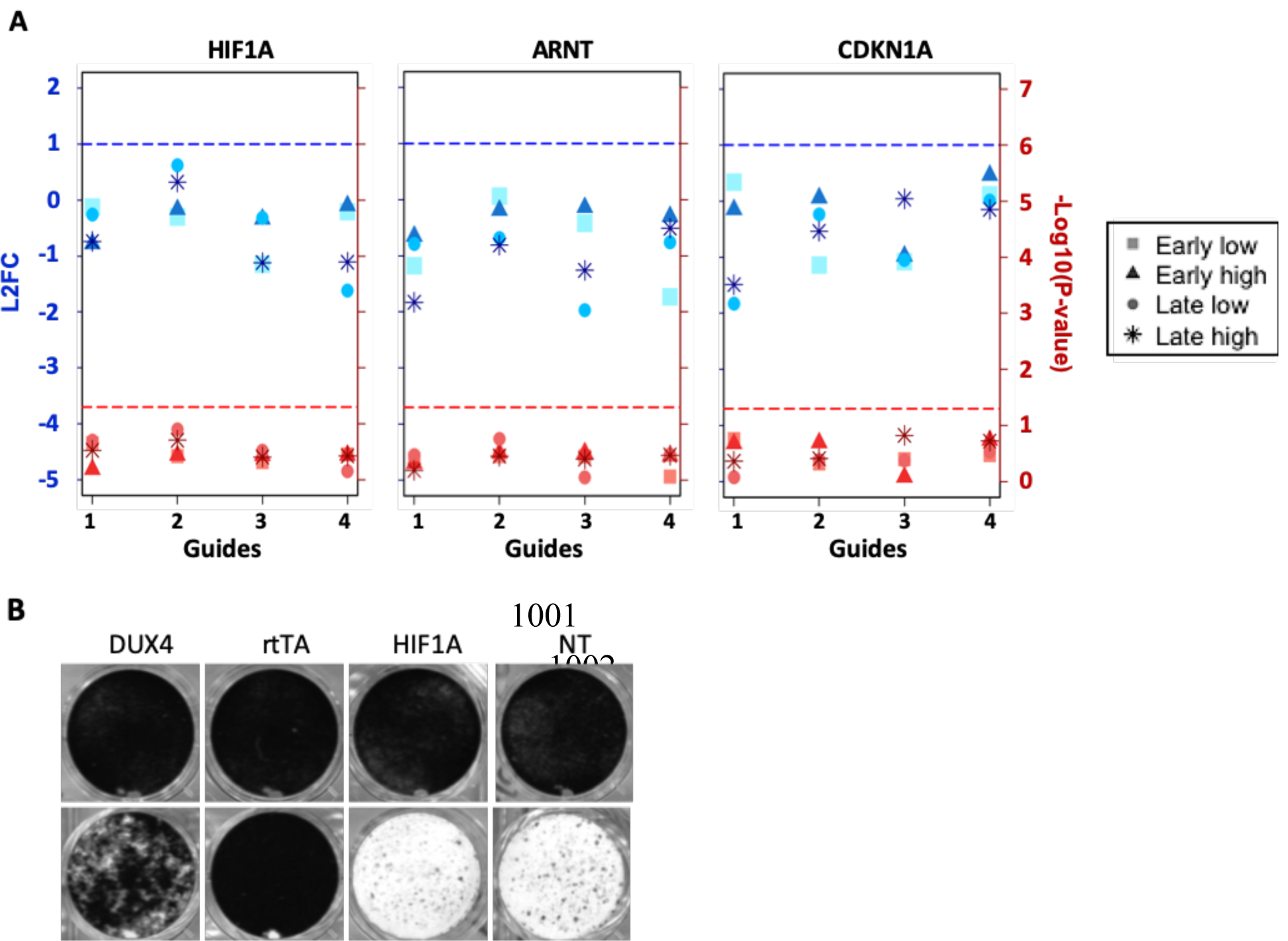
Validation of genes involved in the HIF1 hypoxia pathway. **(A)** Data plots showing the enrichment of sgRNA targeting HIF1A. HIF1B/ARNT, CDKN1A. The LFC value of each individual guide is plotted on the left y-axis, indicated in blue, and the –Log10 P-value is plotted on the right y-axis, in red. Guides located above the blue and red intermitted ablines are considered significant (blue: LFC > 1, red: -Log10 P-value > 1.3). **(B)** Viability staining of DIE-Cas9 uninduced (top panel) and induced with 100ng/ml doxycycline (lower panel) cells, transfected with DUX4, rtTA, HIF1A and non-targeting (NT) sgRNA coding plasmids.

